# Genomic Language Model for Predicting Enhancers and Their Allele-Specific Activity in the Human Genome

**DOI:** 10.1101/2025.03.18.644040

**Authors:** Rekha Sathian, Pratik Dutta, Ferhat Ay, Ramana V. Davuluri

## Abstract

Predicting and deciphering the regulatory logic of enhancers is a challenging problem, due to the intricate sequence features and lack of consistent genetic or epigenetic signatures that can accurately discriminate enhancers from other genomic regions. Recent machine-learning based methods have spotlighted the importance of extracting nucleotide composition of enhancers but failed to learn the sequence context and perform suboptimally. Motivated by advances in genomic language models, we developed DNABERT-Enhancer, a novel enhancer prediction method, by applying DNABERT pre-trained language model on the human genome. We trained two different models, using large collection of enhancers curated from the ENCODE registry of candidate cis-Regulatory Elements. The best fine-tuned model achieved 88.05% accuracy with Matthews correlation coefficient of 76% on independent set aside data. Further, we present the analysis of the predicted enhancers for all chromosomes of the human genome by comparing with the enhancer regions reported in publicly available databases. Finally, we applied DNABERT-Enhancer along with other DNABERT based regulatory genomic region prediction models to predict candidate SNPs with allele-specific enhancer and transcription factor binding activity. The genome-wide enhancer annotations and candidate loss-of-function genetic variants predicted by DNABERT-Enhancer provide valuable resources for genome interpretation in functional and clinical genomics studies.

## INTRODUCTION

Eukaryotic enhancers are cis-acting gene regulatory elements, which enhance transcriptional activation, thereby shaping the characteristics and functions of cells, tissue, and organisms. These 100-1500 bp spanning genomic sequences constitute ∼15% of the non-coding part of the human genome, suggesting they are far more numerous than the approximately 20,000 protein coding genes in the human genome (Pennacchio et al. 2013; Panigrahi and O’Malley 2021). Their functional capability hinges on its architecture, including binding affinities, transcription factor (TF) types, and underlying topology (Kurdistani 2012). Disruption in the ideal function of enhancers due to genetic phenomenon like enhancer hijacking, disruption of Topologically Associating Domain (TAD), epigenetic modulation or structural variants can lead to genetic diseases like congenital disorders, cancers, and common complex diseases, collectively termed as enhanceropathies (Kurdistani 2012; Corradin and Scacheri 2014; Herz et al. 2014; Sur and Taipale 2016). A number of Genome-Wide Association Studies (GWAS) have identified risk variants (Blackwood and Kadonaga 1998), mainly SNPs, in enhancer elements, highlighting their clinical relevance in therapeutic strategies (Kurdistani 2012).

The multifaceted nature of enhancers—their varying lengths, ambiguous boundaries, cell-and state-specific activity, orientation-independent function, and variable distance from associated promoters—complicates their computational identification and functional annotation (Blackwood and Kadonaga 1998). In addition, although general features of enhancers are known, there is no consensus on which features need to be present at what levels to declare a region as “enhancer”. Considering the profound biological and clinical significance of enhancers, the research community consistently endeavored to identify and better understand these non-coding regions (Consortium et al. 2020). While several experimental methods, such as ChIP-seq (Visel et al. 2009), ATAC-seq (Buenrostro et al. 2013), DNase-seq (Boyle et al. 2008), CAGE (Guerrini et al. 2022), GRO-seq (Lopes et al. 2017), PROseq (Kwak et al. 2013) and others (Kwasnieski et al. 2012; Melnikov et al. 2012; Arnold et al. 2013), have been applied for genome-wide mapping of candidate enhancers, their precise identification comprehensively across many conditions and cell types has remained elusive. This led to development of a flurry of ML-based predictive models for enhancer identification.

One of the most widely used approaches is the use of epigenetic information, for example, ChromaGenSVM (Fernandez and Miranda-Saavedra 2012) uses histone marks, CSI ANN (Firpi et al. 2010) utilizes the chromatin signatures and RFECS (Rajagopal et al. 2013) integrates histone modification profiles. These methods often fail to learn the actual context of DNA sequences of the functional region. Later, more efficient methods like iEnhancer-2L (Liu et al. 2016a), iEnhancer-XG (Cai et al. 2021), iEnhancer-EL (Liu et al. 2018) and EnhancerPred (Jia and He 2016) have been developed. These methods focused on extracting sequence features or nucleotide composition, with improved accuracy than the earlier methods. Furthermore, while some recent methods have applied Convolutional Neural network (CNN)-based deep learning approaches (Kim et al. 2016; Liu et al. 2016b; Min et al. 2017; Nguyen et al. 2019), few others have used hybrid methods to leverage the advantages of two different model architectures, for example DEEP (Kleftogiannis et al. 2015), iEnhancer-DCLA (Liao et al. 2022) and BiRen (Yang et al. 2017). More recently, NLP based methods with an additional layer of classification algorithm were developed, like BERT-enhancer (Le et al. 2021) and iEnhancer-ELM (Li et al. 2023). The former method feeds the BERT-based features into a CNN, whereas the latter attaches a 2-layer perceptron network as the classifier to the pre-trained DNABERT model.

Treating DNA sequences like text, we earlier developed DNABERT language model for capturing DNA’s contextual and semantic information in the human genome (Ji et al. 2021), by adapting the Bidirectional Encoder Representations from Transformers (BERT) model that was also applied in many NLP tasks (Lee et al. 2020). DNABERT demonstrated top-tier performance in various downstream tasks including predicting promoters (DNABERT-Prom), splice sites (DNABERT-Splice), and transcription factor (TF) binding regions. An updated version of DNABERT, DNABERT-2 (Zhou et al.) was recently released for multiple species genomes. Based on the performance of DNABERT pre-trained model, we hypothesize that DNABERT can be fine-tuned to precisely predict enhancer sequences, especially focusing on high-attention regions influencing enhancer functionality.

In this study, we curated a large collection of enhancer sequences, which include ENCODE data (Consortium 2012) available through SCREEN (Consortium et al. 2020), and developed a novel enhancer prediction method, called DNABERT-Enhancer. We demonstrated that DNABERT-enhancer captured contextual information from the entire input sequence with attention mechanism achieving superior performance on both independently set aside test data and genome-wide predictions. DNABERT-Enhancer is then applied for predicting functional noncoding SNPs that overlap with the SCREEN enhancers in the human genome. Finally, DNABERT transcription factor fine-tuned models were applied to predict candidate non-coding genomic variants that disrupt TF target regions within the enhancers (**see Figure 1 for brief outline**).

**Figure 1:**
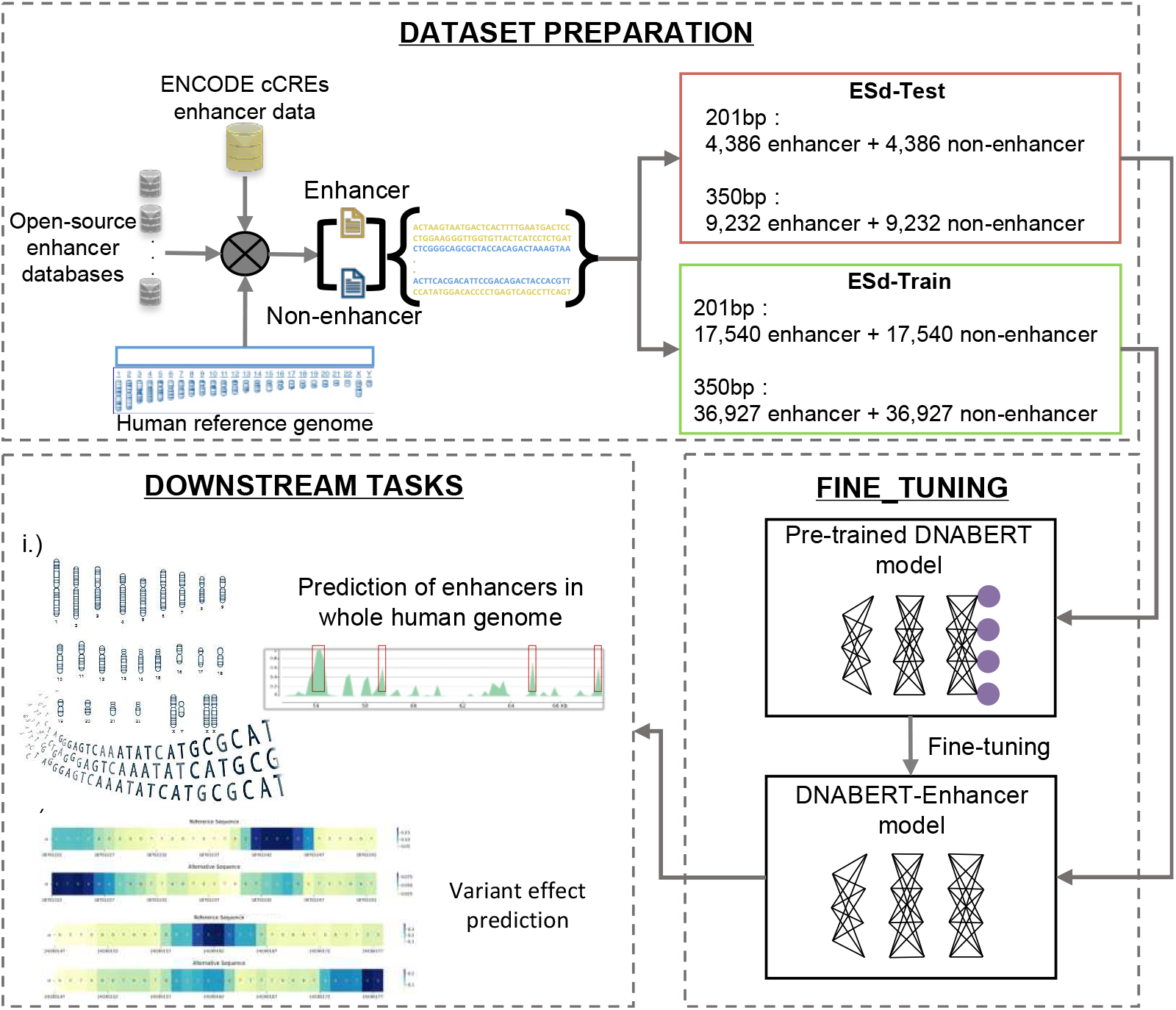
Flowchart illustrating the methods. Enhancer data was initially collected from ENCODE cCREs for creating positive instances. Non-enhancers were created from the human reference genome by eliminating the sequence regions that belong to positive instances along with enhancer data collected from open-source enhancer databases. Pre-trained DNABERT was then fine-tuned with the training data followed by validation. The best model, DNABERT-Enhancer was then used for genome-wide prediction tasks.

## RESULTS

### DNABERT-Enhancer precisely predicts enhancer regions

The datasets used in most published methods have a fixed length of 200bps considering the length of nucleosome and linker DNA. But the catalog of enhancers in different publicly available databases has varying lengths, a few hundreds to more than thousand base pairs. The downloaded registry of cCREs V3, GRCh38 human genome build, have 961,227 enhancers with varying length of 150bp to 350bp. We found two peaks in the length distribution, one at 201bp and the other at 350bp (**Supplementary figure S1**). We, therefore, selected 21,926 (∼2%) enhancers of 201bp length and 46,159 (∼5%) enhancers of 350bp length, as positive instances to fine-tune two different models. The DNABERT-Enhancer-350 model achieved an accuracy of 88.05% with mcc of 76.22%, while the DNABERT-Enhancer-201 model had an accuracy of 82.04% and mcc of 64.27% **(Supplementary** f**igure S3A)**. Hyperparameter tuning played a critical role in refining the performance to maximize the model accuracy while preventing overfitting. The DNABERT-Enhancer demonstrated its best performance at a learning rate of 3e-5, 0.2 warm up rate with 0.0001 weight decay for 350bp length data whereas the model trained on 201bp length data achieved best performance at 0.001 learning rate, 0.1 warm up rate with 0.01 weight decay. **Supplementary figures S1C and S1D** demonstrate the progressive enhancement of our best model’s performance with each step, as evidenced by the increasing trends in accuracy, precision, recall, F1 scores and mcc at minimal train loss. In addition, evaluation based on DNA sequence composition properties (Gupta et al. 2010) showed that the predicted enhancer sequences have higher C and G content, CpG scores and WeSt_Fraction than the non-enhancer sequences, indicating enhancer regions have stronger interactions, C and G triple bonds, compared to the non-enhancer regions (**Supplementary** f**igure S3B-D**).

### DNABERT-Enhancer fine-tuned models outperform baseline classifiers

We validated the performance of DNABERT-Enhancer by comparing the efficiency of both our models to several traditionally used base-line methods namely Random Forest classifier (Breiman 2001), K-nearest neighbor (KNN) (Mucherino 2009), Support Vector Classifier (SVC), Gaussian Naive Bayes, AdaBoost Classifier, and Multi-layer Perceptron (MLP). For the KNN classifier, we utilized the implementation provided by the scikit-learn library, setting the number of neighbors (k) to 3. Compared to deep, transformer-based neural network like BERT which learns high-level representations from raw text, the baseline models often require feature engineering, especially for text data, to learn. For a fair comparison, here we used the embeddings from DNABERT to represent the input sequences thereby facilitating the classification tasks. When validated using ESd-201-Test data, our model outperformed all other base-line methods in terms of all evaluation metrics while with ESd-350-Test data, our model worked best in terms of accuracy, recall, f1 score and mcc (**Table 1**). Though KNN classifier observed to have a higher precision (95.4%), the recall was the lowest (7.64%) across all models indicating that the KNN model overlooks or miss out the true positive values when predicting high confidence enhancer regions. In addition, the mcc of the KNN model was notably low, showing that the algorithm is not working well for the classification task.

**Table 1.**
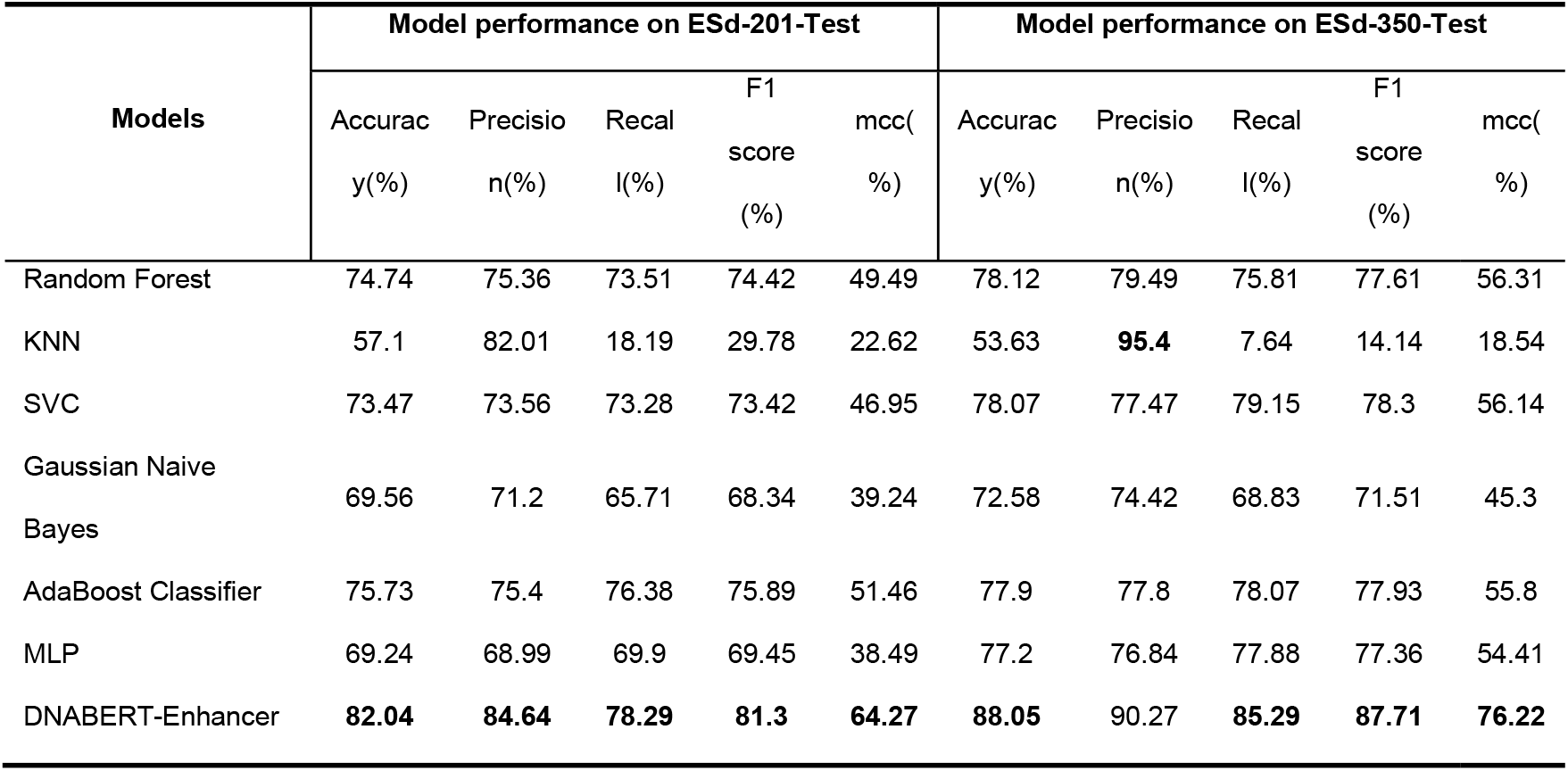
Comparison between DNABERT-Enhancer and the base-line methods. KNN (K-nearest neighbor); Support Vector Classifier (SVC); Multi-layer Perceptron (MLP)

| Models | Model performance on ESd-201-Test |  |  |  |  | Model performance on ESd-350-Test |  |  |  |  |
| --- | --- | --- | --- | --- | --- | --- | --- | --- | --- | --- |
|  | Accurac<br>y(%) | Precisio<br>n(%) | Recal<br>l(%) | F1<br>score<br>(%) | mcc(<br>%) | Accurac<br>y(%) | Precisio<br>n(%) | Recal<br>l(%) | F1<br>score<br>(%) | mcc(<br>%) |
| Random Forest | 74.74 | 75.36 | 73.51 | 74.42 | 49.49 | 78.12 | 79.49 | 75.81 | 77.61 | 56.31 |
| KNN | 57.1 | 82.01 | 18.19 | 29.78 | 22.62 | 53.63 | **95.4** | 7.64 | 14.14 | 18.54 |
| SVC | 73.47 | 73.56 | 73.28 | 73.42 | 46.95 | 78.07 | 77.47 | 79.15 | 78.3 | 56.14 |
| Gaussian Naive<br>Bayes | 69.56 | 71.2 | 65.71 | 68.34 | 39.24 | 72.58 | 74.42 | 68.83 | 71.51 | 45.3 |
| AdaBoost Classifier | 75.73 | 75.4 | 76.38 | 75.89 | 51.46 | 77.9 | 77.8 | 78.07 | 77.93 | 55.8 |
| MLP | 69.24 | 68.99 | 69.9 | 69.45 | 38.49 | 77.2 | 76.84 | 77.88 | 77.36 | 54.41 |
| DNABERT-Enhancer | **82.04** | **84.64** | **78.29** | **81.3** | **64.27** | **88.05** | 90.27 | **85.29** | **87.71** | **76.22** |

### DNABERT-Enhancer outperforms recent enhancer prediction methods

To benchmark DNABERT-Enhancer, we compared its performance to the current state-of-the-art methods, iEnhancer-ELM (Li et al. 2023), iEnhancer-DCLA (Liao et al. 2022) and iEnhancer-ECNN (Nguyen et al. 2019). Our pre-trained DNABERT model was also employed by iEnhancer-ELM model, but a 2-layer perceptron network was attached to the model for classification. Like iEnhancer-ELM, iEnhancer-ECNN is an ensemble method, but the later uses CNN model architecture. iEnhancer-DCLA also utilizes CNN along with Bi-LSTM and attention mechanism to extract features for classification. All of the above-mentioned models were trained on the benchmark dataset from Liu *et al*.(Liu et al. 2018), which contains 1484 enhancers and 1484 non-enhancers constructed based on chromatin state annotations by ChromHMM (Ernst and Kellis 2012).

To avoid any bias due to sequence length, the published methods were tested on both ESd-201-Test and ESd-350-Test datasets. DNABERT-enhancer model outperformed the rest of the methods on ESd-350-Test data in all performance metrics. On ESd-201-Test data, where our model exhibited high performance in all the performance metrics, except specificity (**Supplementary Table ST1**). Though iEnhancer-ELM was observed to have a higher specificity (90.8%) in classifying 201bp sequences, our model showed a significantly improved accuracy and F1 score compared to iEnhancer-ELM (**Figure 2A**). A noteworthy observation was the low mcc score of other models compared to our model. In a binary classification, mcc has proved to be the best evaluation metric, even over F1 score and accuracy as it considers all the four categorical values in a confusion matrix (Chicco and Jurman 2020). Thus, our model has shown superior performance in discriminating enhancers from non-enhancer regions, regardless of the length of the sequences (**Figure 2B and 2C**).

**Figure 2:**
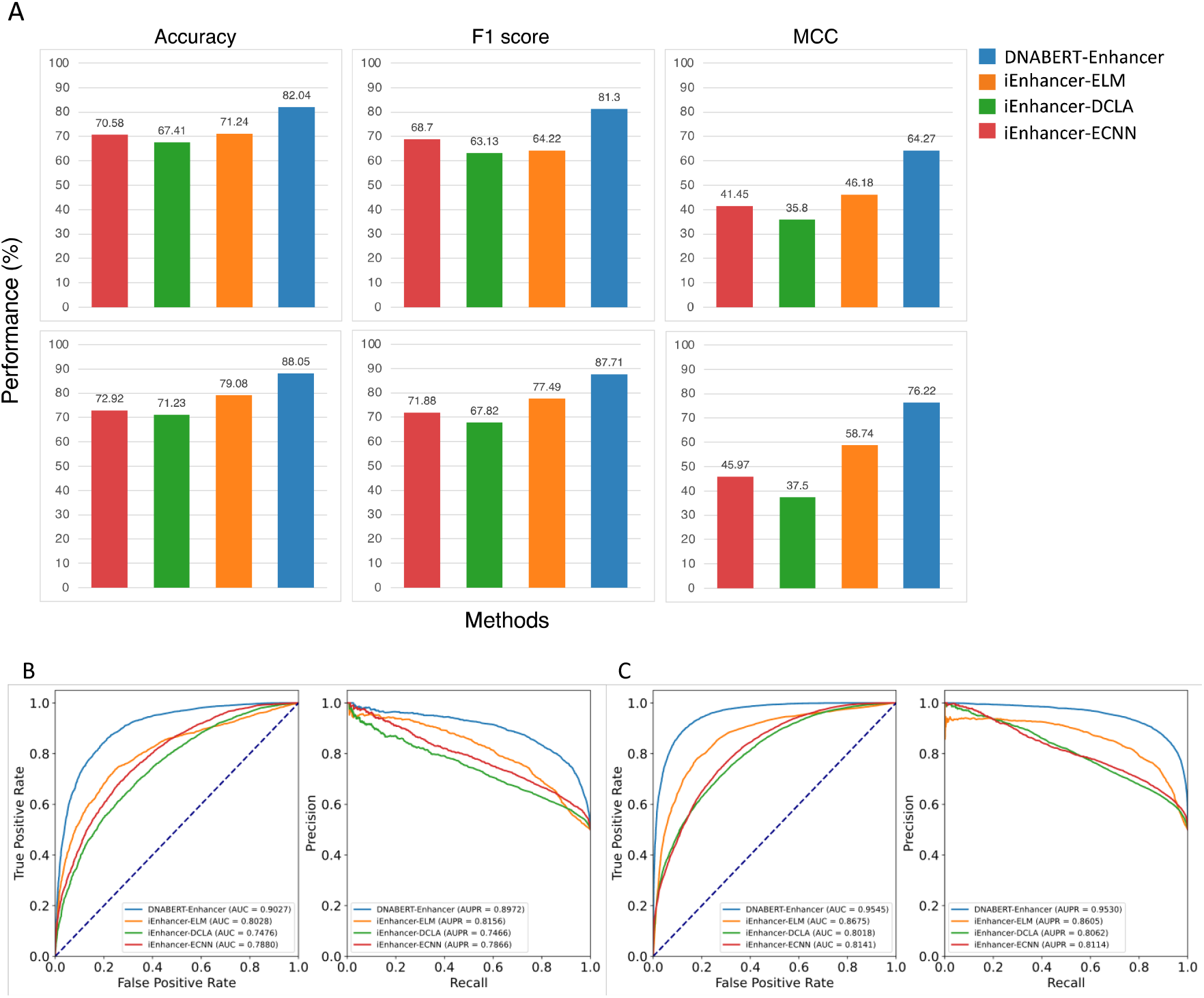
Comparison of DNABERT-Enhancer to the state-of-the-art methods iEnhancer-ELM, iEnhancer-DCLA and iEnhancer-ECNN. (A) Accuracy (left), F1 score (middle) and MCC (right) of the model prediction on ESd-201-Test (top) and ESd-350-Test (bottom). ROC curve and Precision-Recall (PR) curve of the model prediction on (B) ESd-201-Test (top right) and (C) ESd-350-Test (bottom right).

### DNABERT-Enhancer successfully captures the known enhancers in the human genome

The results observed so far showcased the ability of DNABERT-Enhancer in accurately predicting the labeled sequences. However, to evaluate the genome-wide efficacy of our model in capturing the enhancer elements of varying length, we performed prediction on the whole human genome. We applied our best model, DNABERT-Enhancer-350 on a total of 19,582,268 subsequences of 350bp length from the entire human genome. By merging overlapping sequences that were predicted as enhancers (prediction probability ≥ 0.5), we identified a total of 1,822,321 enhancer regions, covering ∼22% of the human genome, including chrX and chrY, in terms of base pairs (**Supplementary Table ST2**). These predicted enhancer regions were further validated by comparing the with the enhancer regions reported in open-source databases.

Since our model was trained on ∼5.6% of the enhancer regions in the ENCODE SCREEN database, we compared the genome-wide prediction to the entire enhancer regions downloaded. About 28.17% of our predicted enhancers mapped to 759,908 enhancer regions in ENCODE SCREEN, which accounts for ∼79.06% of the entire collection (**Figure 3a** and **supplementary Table ST3**). The remaining 21% of the SCREEN collection that were not predicted in the genome-wide analysis are shorter enhancer sequences, with length less than 350bp (**Supplementary figure S4A**). On further analyzing their nucleotide composition, the not predicted ENCODE SCREEN regions observed to have lower fraction of C and G nucleotides, lower CpG content and weaker interactions (A and T double bonds) (**Supplementary figure S4D**), a composition pattern similar to that exhibited by non-enhancer regions. Subsequently we evaluated the remaining DNABERT-Enhancer predicted enhancer regions (∼71%, #1,308,887) which were not present in the SCREEN data using other public data sources for enhancer annotations (see Data sources). We found that 1,300,492 of our 1,308,887 predictions (∼99%; **Supplementary figure S4C**) intersect with enhancers from at least one of these databases, suggesting our methods ability to capture potential enhancers beyond its training data.

**Figure 3:**
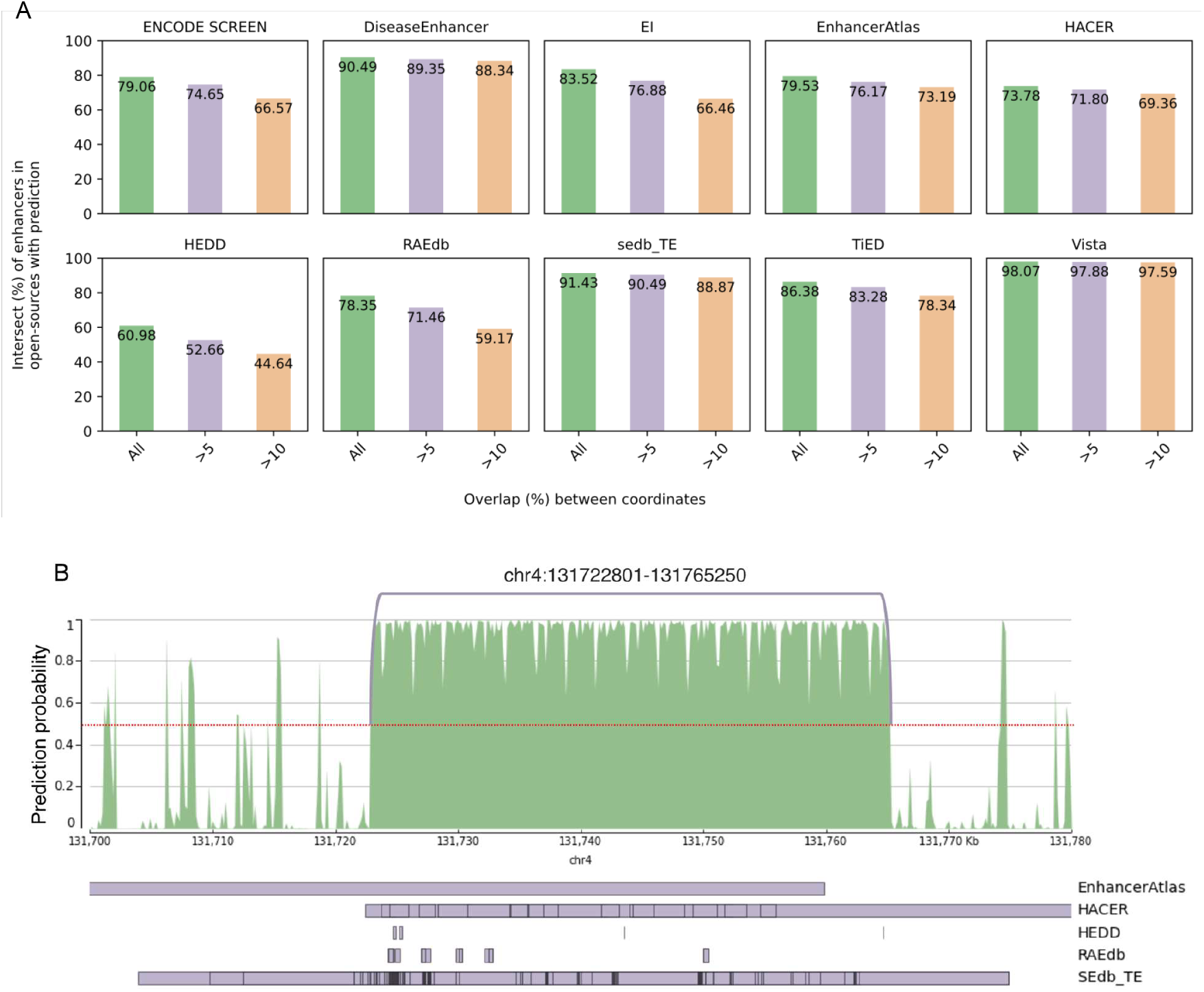
Prediction of DNABERT-Enhancer (350bp) on whole human genome. (A) Intersection of predicted enhancers with enhancers in ENCODE SCREEN as well as 9 different open sources at different intersect overlap cut-offs (>5 and >10). (B) Depicting an enhancer region with length >30kb, predicted by DNABERT-Enhancer, and the overlap of the region with multiple enhancer regions reported in 5 different open-source databases namely EnhancerAtlas, HACER, HEDD, RAEdb and typical enhancers in SEdb.

Further, by comparing the genome-wide prediction results with the publicly available data sources (**Supplementary Table ST4**), we found that the majority of the predicted enhancers were reported in the databases (**Figure 3A**). Notably, ∼91.18% of the predicted enhancers mapped to ∼91.43% of enhancers cataloged in SEdb. Likewise, ∼79.53% of enhancer regions in EnhancerAtlas, an interactive online database of consensus enhancers derived by an unsupervised learning approach using a variety of experimental datasets, intersected with ∼92.80% of the model’s prediction. The database has about 60k enhancers with regions greater than 5 kilobases(kb) and about 1.51M enhancers predicted by our model mapped to these regions, suggesting DNABERT-Enhancer’s ability in capturing elements of super enhancer regions. Furthermore, we found that DNABERT-Enhancer predicted ∼98.07% of the highly confident 1,036 enhancer sequences that were curated by comparative genomics followed by experimental validation in VISTA enhancer browser.

Due to the complicated positioning and orientation of enhancers with respect to their target genes and promoters, identification of a generalized sequence code for enhancers is challenging. The attention regions learned by DNABERT-Enhancer do not have any positional preferences within the enhancer regions (**Supplementary figure S4B**), rather are distributed in different locations of the predicted regions. For the same reason, the overlap of coordinates between the prediction result and the reported enhancers were expected to be partial. Moreover, our approach of merging consecutive predicted enhancer regions identified elements of varying length (**Supplementary Table ST5**) with 4 elements that are larger than 30 kilobases(kb). Interestingly, we observed that these long enhancer regions overlap with different enhancer regions reported in the open-source databases (**Supplementary Table ST6**). **Figure 3B** depicts one such region in chromosome 4 (chr4:131722801-131765250) of length 42,450bp that mapped to varying length enhancer regions reported in 5 different data sources. This evidence further signifies the remarkable capability of DNABERT-Enhancer to recognize and capture the enhancer regions of varying lengths. **Supplementary figure S5** depicts the remaining 3 enhancer regions with length greater than 30kb and their overlap with other enhancer annotation databases.

### DNABERT-Enhancer predicts candidate genetic variant effects in SCREEN enhancer regions

To identify candidate enhancer disruptive variants among the 990 million short variants catalogued in dbSNP database (Sherry et al. 2001), we selected those variants that are located inside SCREEN enhancers of 350bp length. We found that ∼0.52% of the dbSNP variants were mapped to these enhancer regions and repeated the DNABERT-Enhancer-350 prediction model on these sequences with altered alleles. Our analysis identified 62,592 variants exhibiting a disruptive impact on 5,069 SCREEN enhancers of 350bp length (**Supplementary Table ST7**). These enhancer regions were further analyzed for overlapping TF target site disruptions, using DNABERT-TF models that were fine-tuned on the ENCODE ChIP-seq profiles. By using 458 highly accurate DNABERT-TF models (model accuracy is ≥85%), we repeated the predictions on sequences with reference and altered alleles for each of the overlapping dbSNP variants. We identified 15,693 disruptive binding site variants for 349 transcription factors in 3,171 enhancer regions (**Supplementary Table ST8**). Among these, we found a total of 35,447 and 15,146 high-confidence SNPs that were predicted to disrupt enhancers and TF target sites respectively (**Supplementary Tables ST9 and ST10**).

To find variants with known clinical significance, we queried the predicted variants in ClinVar and GWAS databases, leading to the identification of 84 candidate variants with an associated trait or phenotype (**Supplementary Table ST11**). Among those variants, 14 were predicted to disrupt TF binding sites (TFBS) that are located within the enhancer regions. (**Supplementary Table ST12**). Finally, by querying the GTEx catalog of genetic regulatory variants that were predicted to affect gene expression in *cis* across 49 tissues (Consortium 2020), we found 24 variants identified by CAVIAR alone, 83 by DAP-G and 36 by both the methods, totaling 143 high-confidence variants, out of which 56 were predicted to cause disruption of TFBSs.

Here, we present two interesting results (**Figure 4**). An intronic substitution variant mapped to *PCAT19* gene, reported as independent risk variant for prostate cancer in multi-ancestry or ancestry-specific analyses (Wang et al. 2023), was predicted to disrupt MAX (Myc-Associated Factor X) binding site. This variant was reported to influence the expression of *PCAT19* gene in multiple tissues including testis (see **ST12**). Another intronic variant within an enhancer region was predicted to disrupt the ATF4 (Activating Transcription Factor 4) binding site, which was known to be associated with atrial fibrillation mapped to *HS1BP3* gene (see **ST12**). In all the examples, DNABERT consistently shows highest attention at/around the variants of interest. The catalog of predicted variants presented here provide strong candidates for further experimental testing to confirm their functional impact on a specific trait or disease.

**Figure 4:**
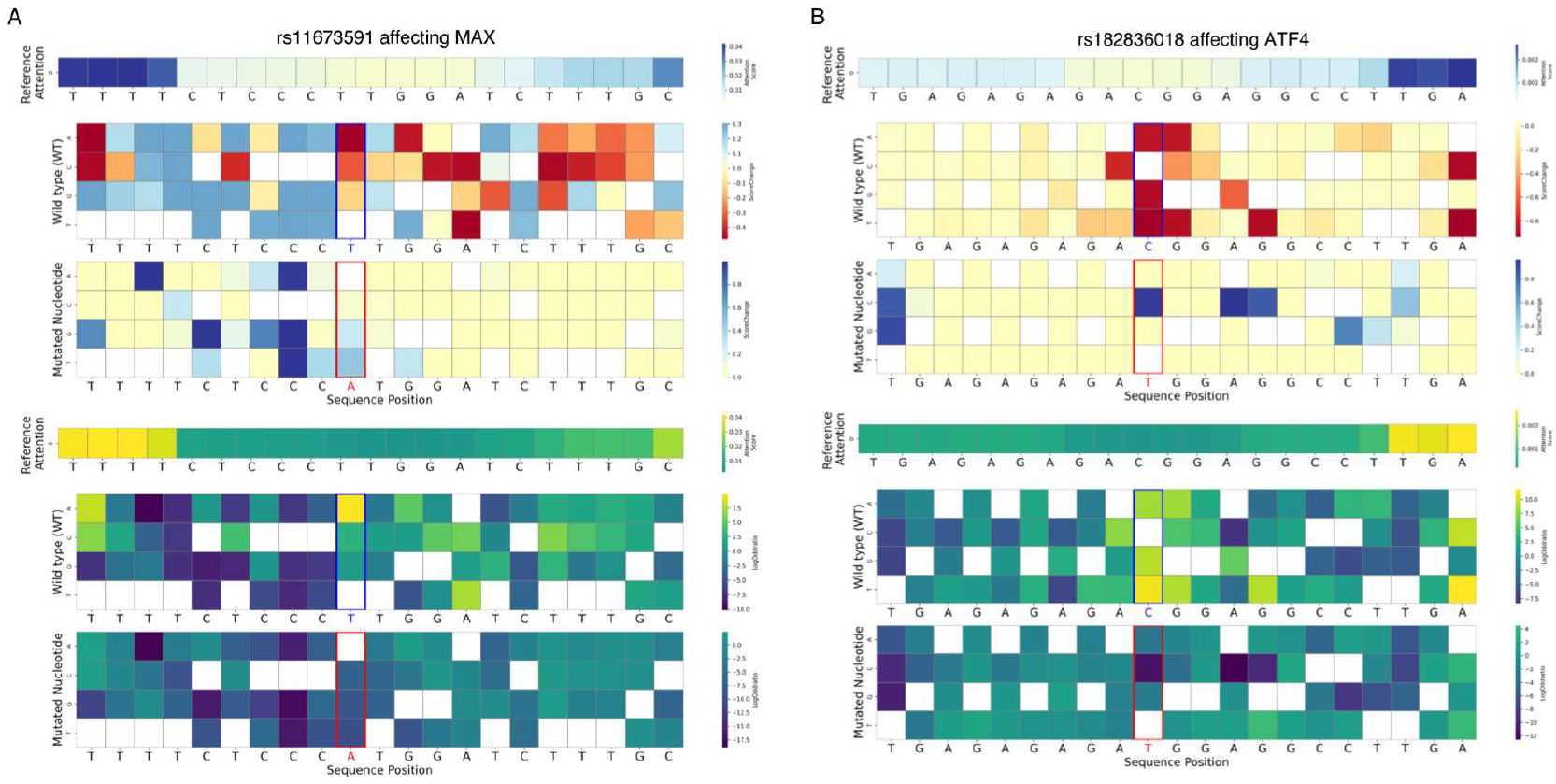
Effect of the functional genomic variants predicted by DNABERT-Enhancer on the TFBS within a given enhancer region. The mutation map depicts the score change (top 3) and the log odds ratio (bottom 3), for wild type and the mutation. (A) an intronic variant affecting MAX binding site targeting PCAT19 gene. (B) an intronic variant in ATF4 binding site targeting HS1Bp3 gene.

## DISCUSSION

Genome-wide prediction of enhancer regions and prioritization of genetic variants that effect the enhancer function are clinically significant and computationally challenging problems (Claringbould and Zaugg 2021). Publicly available databases, such as SCREEN, provide valuable resources for searching candidate regulatory elements, including enhancers, that were derived from ENCODE experimental data (Consortium et al. 2022). However, such experimentally curated enhancer data has limitations due to the finite number of genome-wide epigenetic experiments that can be performed in different cell types and states (Pennacchio et al. 2013). Most of the existing machine-learning based enhancer prediction models were trained on a small dataset, which was curated based on annotations of chromatin-state signatures in 9 cell lines by ChromHMM (Ernst and Kellis 2012). Here, we demonstrated that DNABERT language model can be fine-tuned, by taking advantage of the large collection of enhancer data in SCREEN database. Our results demonstrated that models trained on longer (350bp) sequences outperformed those that were trained on shorter (201bp) sequence data, suggesting longer enhancers contain more regulatory elements and provide more sequence context than shorter regions, an indication of enhancer complexity (Li and Wunderlich 2017). DANBERT-enhancer achieved superior performance in the prediction of enhancer regions in the human genome by largely surpassing existing tools and baseline models. Moreover, DNABERT-enhancer showed remarkable performance in the genome-wide application, where ∼99% of the predicted enhancers overlapped with the enhancer regions recorded in at least one of the surveyed databases. DNABERT-enhancer’s superior performance is attributed to the DNABERT pretrained model, which efficiently captures k-mer based intricate and discriminative language patterns between enhancer regions and other genomic regions by leveraging large number of experimentally curated enhancer sequence data.

GWAS studies, in which millions of SNPs were tested to identify significant genotype-phenotype associations, have revolutionized the field of complex disease genetics. While dbSNP and other related databases provide functional annotation of SNPs located in the protein coding regions, such annotations are mostly lacking for noncoding variants due to lack of understanding of DNA language in the non-coding regions. Moreover, unraveling the causal relationships between specific noncoding genetic variants and diseases remain challenging, due to lack of insights into the underlying molecular mechanisms (Yang et al. 2022; Yang et al. 2024). Here, we demonstrated that DNABERT-enhancer can be applied to directly predict candidate SNPs that are disruptive of enhancers and TF target regions that overlap with the enhancer regions. In summary, we anticipate that DNABERT-enhancer model, released as part of this study, can also be applied to predict enhancers in rodents and non-human primate genomes, without the need for separate fine-training. Furthermore, this approach allows prioritization of potential causal variants resulting in alteration of enhancer function providing candidates for downstream experimental and functional studies, with potential for identification of candidate enhancers for novel gene and cell therapies (Maeder and Gersbach 2016).

## METHODS

### Data sources

To build the fine-tuned models, we collected enhancers available in the Registry of candidate cis-Regulatory Elements (cCREs), which is the integrative layer of the ENCODE Encyclopedia. The cCREs are categorized as regulatory elements based on H3K27ac, H3K4me3 and CTCF signals across 25 human cell types having complete cell-type specific classification and 839 human cell types having partial classification. We also gathered enhancers regions reported in 10 different databases, dbSUPER (Khan and Zhang 2016), DiseaseEnhancer (Zhang et al. 2018), EI cENHs (Pennacchio et al. 2007), EnhancerAtlas (Gao and Qian 2020), HACER (Wang et al. 2019), HEDD (Wang et al. 2018), RAEdb (Cai et al. 2019), sedb (Jiang et al. 2019), TiED (Xiong et al. 2018) and VISTA (Visel et al. 2007) to assist with generating negative instances. Briefly, these databases consist of enhancer sequences identified either through different experimental approaches like CAGE, GRO-seq, PRO-seq, STARR-seq, MPRA or by manual curation. The data was ensured to be in GRCh38 human genome assembly by applying build conversion tools, CrossMap (Zhao et al. 2014) program for hg19 assembly and UCSC LiftOver (Hinrichs et al. 2006) for hg17 assembly. Sequence similarity was also taken care of by excluding sequences with greater than 80% identity.

The negative instances (Non enhancer sequence set) were created from the human reference genome (GRCh38 assembly), wherein regions other than the enhancer regions were designated as non-enhancer regions, resulting in a total of ∼6,8750,000 non-enhancer sequences. Like enhancers, the non-enhancer sequences were also fragmented into a fixed length of 201bp and 350bp, thereby creating two different datasets of negative instances as well. The positive and the negative instances thus created had a huge disparity in terms of the size of the dataset. To avoid the model from exhibiting bias towards a specific class or skewed outcomes of performance metrics, we balanced the dataset by 1:1 enhancer to non-enhancer ratio.

### Fine-tuning of DNABERT pre-trained model

#### Training and Testing of DNABERT-enhancer models

We built two different fine-tuned models – DNABERT-Enhancer-201 and DNABERT-Enhancer-350 for predicting enhancer sequences of length 201 bp and 350 bp respectively. These models were trained on 80% of enhancer sequences – (ESd-201-Train, ∼17,500 enhancers & ∼17,500 non-enhancers randomly selected from each class; and ESd-350-Train, ∼36,900 enhancers & ∼36,900 non-enhancers randomly selected from each class). The best trained models were tested on the remaining 20% of the sequences that were held out (ESd-201-Test, ∼4,300 each in enhancer and non-enhancer classes and ESd-350-Test, ∼9,200 each in enhancer and non-enhancer classes). To optimize the model, we fine-tuned the hyperparameters: learning rates ranging from 1e-3 to 3e-6, warm-up steps set at 0.1, and 0.2, and weight decay as 0.0001, 0.001, 0.005, 0.01 and 0.02, maintaining a consistent dropout throughout. The fine-tuning process ran on 8 NVIDIA A40 GPUs for approximately 4-5 hours for one set of hyperparameters. In **supplementary figures-S1B-D**, we provide further details of all the performance metrics and loss patterns of our best model during training phases.

#### Application of DNABERT-enhancer models on the human

To validate the efficiency of the model in genome-wide analysis, we downloaded the human genome sequence data (GRCh38) from UCSC genome browser. Each chromosome was processed to create subsequences of length 350bp, by applying a sliding window of 150bp. Sequences containing ‘N’ nucleotides were dropped from the data as they are unidentified bases within the sequences. The data was then tokenized to hexamers (6-mer), followed by random labeling, and were given to the best fine-tuned enhancer model for the prediction. Once the prediction results were generated for subsequences (*S*_*i*_|*i* ∈ [1,2,3 … *m*]), where *m* is the total number of subsequences, we deduced the probability score *P*(*W*_*j*_) for every 150bp window (*W*_*j*_|*j* ∈ [1,2,3 … *n*]), where *n* is the total number of windows. The window was labeled as an enhancer if its probability *P*(*W*_*j*_) ≥ 0.5. And if two or more consecutive windows had probability, say *P*(*W*_*j*_) ≥ 0.5 and *P*(*W*_*j*+1_) ≥ 0.5, then the regions were merged into a single continuous enhancer prediction (**Supplementary figure S2**). The predicted enhancer regions were then evaluated by overlapping with the enhancer data cataloged in the open-source databases, at different overlap cutoffs (in percentage; 5,10,15,20,25,50,75,95 and All).

### Variant effect prediction

To predict the effect of genetic variants within the enhancers and the TF target regions, we downloaded the dbSNP release 155(GRCh38) (Sherry et al. 1999) and surveyed for the short variants that fall in those regions. The prediction probabilities were then recomputed by substituting the variant nucleotide change in the sequence. To access the effect of variant in the TF target regions within the enhancers, we followed the same procedure but using the DNABERT finetuned models on TFs built from ChIP-seq peaks (enriched genomic regions) downloaded from the ENCODE portal. Further, the effect was measured by calculating the score change and log odds ratio as given in (Ji et al. 2021). Briefly, score change is the difference between the probabilities, defined as *S* = Δ*p* = (*p*(*s*^’^) − *p*(*s*))*max*(*p*(*s*^’^), *p*(*s*)), where *p*(*s*) is the prediction of sequence, *p*(*s*^’^) is the prediction of substituted sequence and the max term is added to amplify the strong effects of certain genetic variants. The Log odds ratio shows the association of two events, calculated by 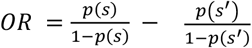. A log odds ratio of 0 indicates no association between the events. To predict candidate variants that have a disruptive functional effect on enhancer sites, we downloaded the dbSNP_clinvar155 track, ClinVar (Landrum et al. 2018) and GWAS catalogue (Uffelmann et al. 2023). These databases contain comprehensive information on the clinical and functional aspects of many variants. In addition, the SNPs were analyzed further for the expression quantitative trait loci (eQTL) using the UCSC tracks for high-confidence cis-eQTLs from CAVIAR and DAP-G displaying gene/variant pairs for 49 GTEx tissues (V8 data release), which are located within 1MB of gene transcription start sites. CAVIAR confidence set consist of 526,252 unique high causal variants with causal posterior probability > 0.1, whereas DAP-G track consist of 1,451,916 unique variants with strong eQTLs signals, signal-level posterior inclusion probability (SPIP) > 0.95, totaling to 1,674,664 variants.

### Evaluation metrics

The performance of the fine-tuning tasks was measured by 6 performance metrics: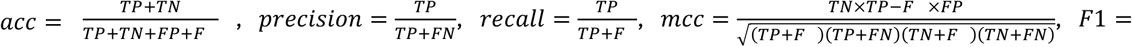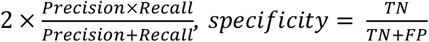, where TP is true positive, TN is true negative, FP is false positive, and FN is the false negative values.

### Sequence properties

To study the nucleotide composition of DNA sequences, we applied the sequence composition-based properties in (Gupta et al. 2010). Briefly, a total of 10 properties were studied. Properties based on single nucleotide were A_fraction, C_Fraction, G_Fraction, T_Fraction, PurPyr_Fraction (fraction of purines and pyrimidines), AmKe_Fraction (fraction of amino bases and keto bases) and WeSt_Fraction (fraction of weak and strong bases). Remaining 3 properties were to assess the CpG islands. Sequence property CpG1 was based on di-nucleotide composition whereas properties CpG2 and CpG3 were based on tri-nucleotide composition. All calculations were based on the equations stated in the original article.

## DATA ACCESS

All code and datasets supporting the findings of this study are publicly available in the GitHub repository **DNABERT-Enhancer** at https://github.com/DavuluriLab/DNABERT-Enhancer. Detailed instructions for reproducing the results are provided within the repository.

## ACKNOWLEDGEMENTS

We thank all members of the Davuluri lab (The State University of New York at Stony Brook) and Ferhat Ay (La Jolla Institute for Immunology) for critical discussions and helpful advice. This work was financially supported by grants from National Library of Medicine/National Institutes of Health funding – [R01LM01372201 to R.D., R35GM128938 to F.A]

## AUTHOR CONTRIBUTIONS

Rekha Sathian: Conceptualization [lead], Data curation [lead], Formal Analysis [equal], Investigation [equal], Methodology [equal], Resources [equal], Software [equal], Validation [equal], Visualization [lead], Writing—original draft [lead]. Pratik Dutta: Formal analysis [equal], Investigation [supporting], Methodology [equal], Resources [supporting], Software [supporting], Validation [supporting], Visualization [supporting], Writing—review & editing [supporting]. Ferhat Ay: Writing – review & editing [supporting]. Ramana V Davuluri: Conceptualization [equal], Funding acquisition [lead], Project administration [lead], Supervision [lead], Writing—original draft [supporting], Writing—review & editing [supporting].

